# Human tissues exhibit diverse composition of translation machinery

**DOI:** 10.1101/2023.01.16.524297

**Authors:** Aleksandra S. Anisimova, Natalia M. Kolyupanova, Nadezhda E. Makarova, Artyom A. Egorov, Ivan V. Kulakovskiy, Sergey E. Dmitriev

## Abstract

While protein synthesis is vital for the majority of cell types of the human body, diversely differentiated cells require specific translation regulation. This suggests specialization of translation machinery across tissues and organs. Using transcriptomic data from GTEx, FANTOM, and Gene Atlas we systematically explored the abundance of transcripts encoding translation factors and aminoacyl-tRNA synthetases (ARSases) in human tissues. We revised a few known and identified several novel translation-related genes exhibiting strict tissue-specific expression. The proteins they encode include eEF1A1, eEF1A2, PABPC1L, PABPC3, eIF1B, eIF4E1B, eIF4ENIF1, and eIF5AL1. Furthermore, our analysis revealed a pervasive tissue-specific relative abundance of translation machinery components (e.g. PABP and eRF3 paralogs, eIF2B subunits, eIF5MPs, and some ARSases), suggesting presumptive variance in the composition of translation initiation, elongation, and termination complexes. These conclusions were largely confirmed by the analysis of proteomic data. Finally, we paid attention to sexual dimorphism in the repertoire of translation factors encoded in sex chromosomes (eIF1A, eIF2γ, and DDX3), and identified testis and brain as organs with the most diverged expression of translation-associated genes.

## 1 Introduction

In multicellular organisms, differences between cell types are largely defined by specific patterns of gene expression. The cell specialization is partially determined at the translational level. Multiple studies showed that gene expression profiles at the level of transcription and translation vary across organs (see [1,2] and references therein). However, the differential regulation of protein synthesis across cell types and tissues is still poorly studied [3,4].

Recent advances in high-throughput approaches gave a new impetus to research in this field [5]. Cell type-specific expression of tagged ribosomal proteins provided a methodology for affinity purification of polysomal mRNAs from defined cell populations within a complex tissue [6–8]. These approaches were further enhanced by integration with ribosome profiling (Ribo-Seq), a technique that allows the analysis of ribosome occupancy of mRNA on a transcriptome-wide scale [9,10]. Although these techniques enabled quantitative and qualitative characterization of translatome from specific cells and lineages [10,11], existing studies have a limited ability to assess a *bona fide* translation efficiency of particular transcripts due to a lack of the complementing transcriptome information (RNA-Seq) from the same cells. Recently, studies involving Ribo-Seq of animal tissues and organs revealed organ-specific differences in translation efficiency and elongation speed [12–15]. Other findings revealed different tRNA repertoires and codon usage in different tissues [16,17], as well as peculiar features like readthrough rate [18]. However, to which extent the tissue-specificity of gene expression is determined at the translation level is still a poorly resolved issue.

It is clear that the variability of the transcriptome across cell types requires specialization of the translation apparatus. Although protein synthesis machinery is indispensable for life and hence thought to be ubiquitously present, recent systematical analyses revealed that some ribosomal components [19–21] and tRNAs [22,23] demonstrate tissue-specific expression. This is likely true for other components of translation machinery, such as translation factors, aminoacyl-tRNA synthetases (ARSases), and other auxiliary proteins, but only a few cases of tissue-specific expression of these components were described until now.

The classic example is the well-documented tissue-specific expression of two eEF1A paralogs in rodents, with eEF1A2 present only in brain, heart, and muscle, and eEF1A1 expressed in all other tissues [24,25]. Testis-restricted transcription was reported for the human *PABPC3* gene [26], which is absent in mice, while gonad-specific expression of other poly(A)-binding protein paralog, PABPC1L/ePAB, is controlled at both transcriptional and post-transcriptional levels in mice and humans [27,28]. An elevated concentration in testis was also shown for translation elongation factor eIF5A2 [29]. Diverged cap-binding protein eIF4E1B was declared to be oocyte-specific in some vertebrates (for review, see [30]), although its expression has not been extensively analyzed across multiple tissues and organs.

The above fragmentary information indicates that the expression of selected translation machinery components is restricted to particular tissues or cell types. It is even more intriguing, that abundance of canonical translation factors, like eIF2γ, eIF5, ETF1/eRF1, GSPT2/eRF3b, eEF1B subunits, or mitochondrial translation factors, also varies significantly across tissues [31–40]. Thus, the uniformity of translation apparatus in different cell types is illusory, suggesting that distinct translational properties of human cell types exist that provide an additional layer of tissue-specific regulation of gene expression.

As mammalian tissues require diverse levels of protein synthesis activity [41–43], it is expected that some of the tissues have more translation machinery components, including ribosomes, than the others [19,20,44]. However, it remains unclear whether this difference significantly affects translation regulation, as in many cases it is determined not by the absolute but by the relative abundance of translation factors [45]. To our knowledge, no such analyses have been performed yet.

Here, using gene expression data from multiple sources, we systematically analyzed the abundance of mRNAs encoding translation factors and ARSases across human tissues. We found intriguing differences in the transcript levels, not only in terms of the absolute values but also in relative shares within physically and functionally related complexes. The diverged expression repertoire of translation-associated genes was especially prominent in testis and brain. Where possible, we confirmed these observations by analysis of proteomic data.

## 2 Methods

Data analysis and visualization were done in R environment. We analyzed mRNA abundance data of different tissues and primary cells available in FANTOM5 (http://fantom.gsc.riken.jp/5/), GTEx (https://gtexportal.org/), and Gene Atlas [46] databases, while mass-spectrometry data were taken from [1]. We considered the genes encoding protein synthesis machinery components according to Table 1.

**Table 1.**
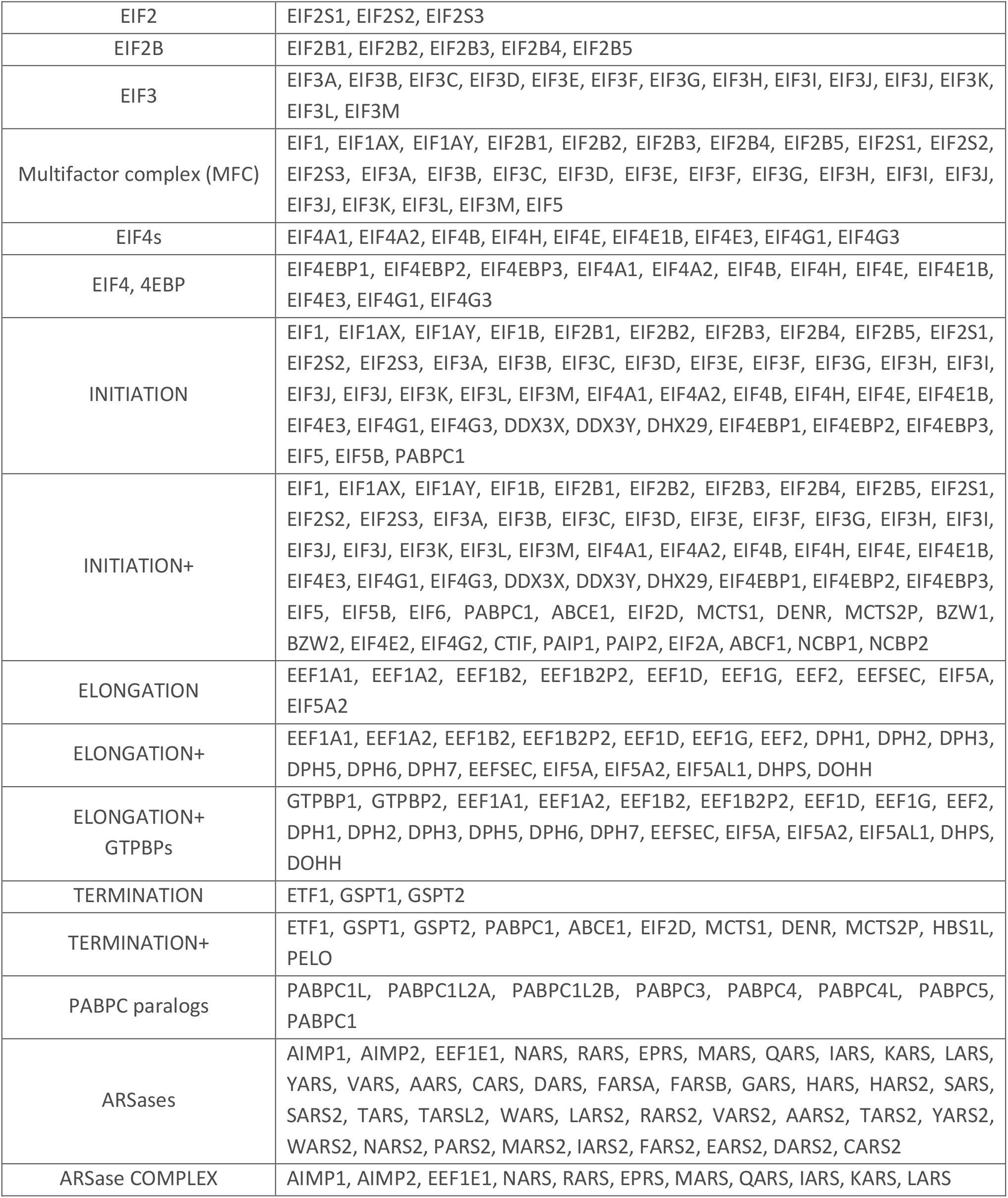
Complexes and functional groups of translation-associated proteins classified according to functional and physical interactions.

The CAGE data (cap analysis of gene expression) from FANTOM5 (http://fantom.gsc.riken.jp/5/) phase 1 mapped to hg19 genome assembly were downloaded as normalized TPM (Tags Per Million) for each transcript, as provided in the FANTOM5 data, and summed up across transcripts to obtain the gene-level expression estimates. Samples with extreme normalization factors (less than 0.7 or higher than 1.4) were excluded from the analysis. The Gene Atlas data [46] were obtained from the BioGPS portal (http://biogps.org/) as normalized expression units from the Human U133A/GNF1H microarray. Probe sets represented in U133A annotation were selected for further analysis, and the mean expression across probes was considered as the estimate of gene expression. GTEx v7 RNASeQCv1.1.8 gene expression data were downloaded as TPM (Transcripts Per Kilobase Million) from https://gtexportal.org/home/datasets.

In our analysis, we used absolute gene expression estimates as well as relative expression estimated as a share of a particular gene in the total gene expression of a particular complex (see Table 1). Relative estimates were used to analyze the target gene expression in the context of a protein complex with a particular functional role. For the latter analysis, we classified the gene complexes according to the functional and physical interactions of the respective proteins (Table 1). The use of relative gene expression allowed us to address the changes in relative transcript abundance in the context of the expression of protein partners and the stoichiometry of the complexes.

To obtain a reliable statistical estimate of the tissue-specific contribution of a particular protein to a particular complex or group, we used GTEx data and the following approach. First, for each complex, we excluded the samples with the total expression of genes of the complex less than half a mean across samples. Next, for each member of the complex, we estimated its share in the total expression of the complex. With the vector of shares across samples, we performed set enrichment analysis on the ranked list of samples. To this end, we classified the samples according to the tissue of origin, e.g. ‘brain’ (30 groups of 11688 samples in total according to GTEx metadata), and checked the positions of samples of a particular group in the total ranked list of samples. Normalized enrichment scores (NES) and statistical significance were estimated with fgsea R package [47] (10000 permutations). P-values were corrected for multiple testing using FDR correction for the number of sample groups.

## 3 Results

### 3.1 Characterization of a tissue-specific expression pattern of two human eEF1A paralogs using an integrated transcriptomic and proteomic data analysis

We analyzed transcriptomic data from FANTOM, GTEx, and Gene Atlas databases, available for various human tissues, to assess differential expression of genes encoding translation factors, ARSases, and auxiliary proteins (hereafter called TAG, for translation-associated genes). Since different cell types clearly have distinct requirements for protein synthesis and its efficiency, expected levels of translation machinery components vary in a wide range. Therefore, to access the differences in the composition of translation machinery across tissues, we analyzed the relative abundance of the TAG transcripts within functional complexes. To this end, we determined the number of complexes and functional groups of translation-associated proteins (Table 1). TAGs were combined into several groups based on known physical and functional interactions of their products. Each factor could be included in more than one group, and selected smaller groups were fully included in the larger groups. Next, we calculated two parameters for each gene, the transcript abundance value (see Materials and Methods), and the relative abundance of the transcript within the group. The latter allowed us to detect putative differences in the stoichiometry of the complexes.

The classic example of TAGs with a highly pronounced tissue-specific expression is the genes encoding rodent eEF1A. The translation elongation factor eEF1A delivers aminoacyl-tRNA to the ribosomal A-site for decoding, which is inevitable for protein synthesis, and is one of the most abundant proteins in the cell [48]. In many eukaryotes, eEF1A is encoded by two paralogous genes. In mice and rats, these genes (*Eef1a1* and *Eef1a2*) have a well-documented mutually exclusive expression pattern [25,49]. Here, we used the example of their human orthologs, *EEF1A1*, and *EEF1A2*, to illustrate a strategy for the characterization of tissue-specific TAG expression.

Analysis of GTEx data showed that human *EEF1A2* is exclusively expressed not only in brain, heart, and muscle (similarly to the rodent gene), but also in the pituitary and, to less extent, in adrenal and salivary glands (Figure 1A). The expression pattern of *EEF1A2* was in agreement with proteomics data from one of the most representative proteomics atlas of human tissues (Figure 1B) [1]. Notable, the gland-specific expression of this gene has not been documented before. As expected, the expression of *EEF1A1* mirrored that of *EEF1A2*, being reduced in the organs and tissues with elevated *EEF1A2* expression (Figure 1A,B).

**Figure 1.**
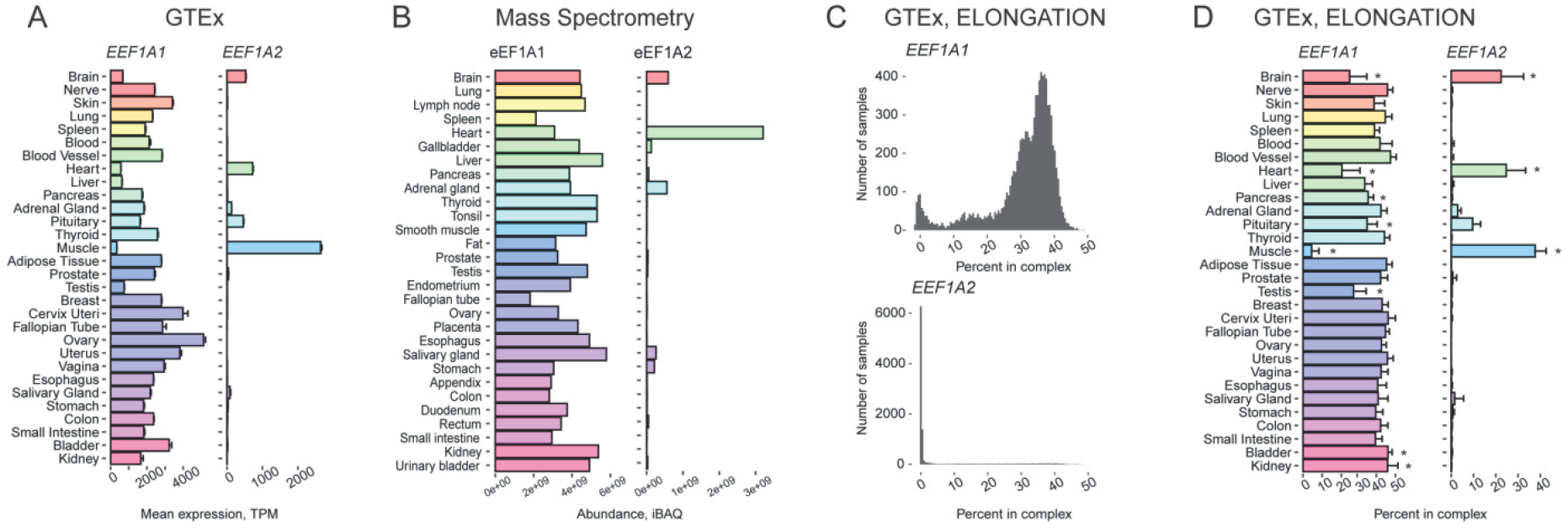
Tissue-specific expression pattern of two human genes encoding eEF1A paralogs, *EEF1A1* and *EEF1A2*. **(A)** Expression of the *EEF1A1* and *EEF1A2* genes in various human tissues according to GTEx. **(B)** Levels of eEF1A1 and eEF1A2 proteins in various human tissues according to high-throughput proteomic analysis [1]. **(C)** Histograms showing the distribution of the percentage of *EEF1A1* (top) and *EEF1A2* (bottom) expression among the genes from the “ELONGATION” complex across various human tissues, according to GTEx. **(D)** Percentage of *EEF1A1* and *EEF1A2* expression among the genes from the “ELONGATION” complex across various human tissues, according to GTEx. TPM, Transcripts Per Kilobase Million; *, FDR corrected p-value < 0.01 in enrichment analysis (fgsea R package).

Interestingly, analysis of eEF1A1 mRNA and protein representation among components of the ELONGATION complex revealed that, while its relative abundance was reduced in skeletal muscle in comparison with most other tissues, in some brain samples it remained at a high level, indicating the simultaneous presence of two eEF1A paralogs in these samples (Figure 1 C,D). Similar results were obtained with data from Gene Atlas and FANTOM databases (Supplementary Figure 1 A,B). Due to a slightly different set of organs and tissues analyzed in the three projects, expression in tongue, eye, throat, and pineal gland also became evident.

### 3.2 Tissue specificity of translation-associated proteins encoded in sex chromosomes

To further validate our approach, we assessed the tissue specificity of translation-associated proteins encoded by sex chromosomes. Obviously, the expression of Y-encoded TAGs should not be detected in female-specific tissues. Among 65 protein-encoding genes located at the human Y chromosome [50] there are only two TAGs from our list, *EIF1AY* and *DDX3Y*. Both of them have paralogs at the X chromosome, *EIF1AX* and *DDX3X*, respectively.

*EIF1AX* and *EIF1AY* are paralogs coding for the canonical translation initiation factor eIF1A, which is indispensable for 48S preinitiation complex formation [45]. Two eIF1A variants are almost identical (differ by only 1 amino acid out of 144, Leu50 in eIF1AY and Met50 in eIF1AX in humans) and thus probably have identical functions. An equal overall eIF1A level in male and female cells thus requires that the *EIF1AX* gene escapes X chromosome inactivation, which is indeed the case [51,52]. Analysis of *EIF1AX* and *EIF1AY* mRNA abundance confirmed the absence of the *EIF1AY* transcript in female-specific organs and revealed differential expression between tissues (Supplementary Figure 2A). We then calculated their relative abundance within the group “INITIATION”. This approach revealed a slightly variable relative abundance of the *EIF1AX* mRNA, while the expression level of the *EIF1AY* gene was not uniform (Figure 2A). This analysis also showed a predominance of the *EIF1AX* gene as a source of the eIF1A protein but pointed to the heart as an organ with an apparently higher overall eIF1A level due to the elevated impact of *EIF1AY* expression. The proteomic data [1] available only for eIF1AX revealed its depletion in testis in comparison with other factors of the INITIATION complex (Supplementary Figure 2B). This was not seen in transcriptomic data from either source and suggested an additional layer of regulation at the post-transcriptional level, as well as a putative role in specialized translational control in this organ. Interestingly, the presence of the Y-encoded eIF1A isoform is not a conserved feature in mammals, as mice have only one gene, *Eif1ax*, which does not escape X inactivation [53].

**Figure 2.**
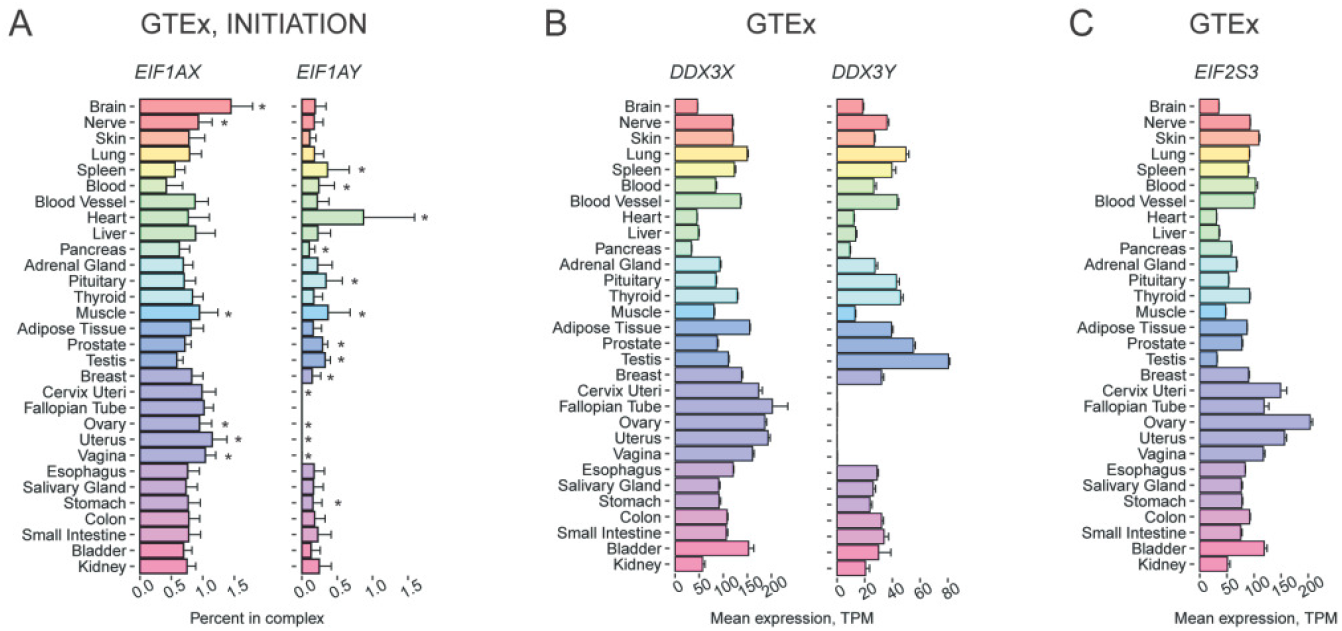
Tissue-specific expression pattern of translation-associated genes localized in sex chromosomes. **(A)** Percentage of *EIF1AX* and *EIF1AY* expression among the genes from the “INITIATION” complex across various human tissues, according to GTEx. **(B)** Expression of the *DDX3X* and *DDX3Y* genes in various human tissues according to GTEx. **(C)** Expression of the *EIF2S3* gene in various human tissues according to GTEx. TPM, Transcripts Per Kilobase Million; *, FDR corrected p-value < 0.01 in enrichment analysis (fgsea R package).

*DDX3X* and *DDX3Y* produce isoforms of DEAD-box helicase, DDX3, an auxiliary translation initiation component, which plays an essential role in eukaryotic RNA metabolism [54]. The two proteins have only 92% identity. *DDX3X* cannot substitute for *DDX3Y* function, while the replacement of *DDX3X* with *DDX3Y* does not affect the translation rate [55–57]. Deletions encompassing the *DDX3Y* gene are thought to result in spermatogenic failure (reviewed in [58,59]), although this opinion has been challenged recently [60]. Mutations in *DDX3X* are a cause of intellectual disability in females and a decreased viability in males (reviewed in [61]), which, as Venkataramanan et al. suggest, can be explained by tissue-specific expression of *DDX3X* and *DDX3Y* [57]. Similar to *EIF1AX*, *DDX3X* escapes X-chromosome inactivation [53].

Our data analysis showed the absence of *DDX3Y* mRNA and protein in female-specific tissues that met our expectations. In contrast, the protein level was high in testis. *DDX3X* is expressed ubiquitously; however, the abundance of its transcript is somewhat higher in the female reproductive system (Figure 2B), but this slight difference is not reflected in proteomics data (Supplementary Figure 2C).

In humans, X-chromosome contains one more unevenly expressed TAG – *EIF2S3*. It codes for the γ-subunit of eIF2, a critical translation machinery component required for Met-tRNAi delivery during initiation complex formation. GTEx data analysis revealed that the *EIF2S3* transcript abundance is higher in female reproductive tissues than in testes (Figure 2C), which is in agreement with its X-inactivation escape profile [62]. The relative expression of eIF2 complex components (Supplementary Figure 2D) varies between female and male reproductive tissues accordingly. Interestingly, mice have a paralog of *Eif2s3x, Eif2s3y*, which is required to drive spermatogenesis and represents one of the two genes irreplaceable for male mice fertility [63]. Mouse *Eif2s3x* escapes from X inactivation similarly to the human gene [53,62] and is expressed higher in female mice than in males in developing and adult brains at the transcriptional level [31]. Notably, mutations in the human *EIF2S3* result in MEHMO syndrome (Mental retardation, Epileptic seizures, Hypogenitalism, Microcephaly, and Obesity) in males [64].

### 3.3 A number of translation factors have a pronounced tissue-specific representation within corresponding functional groups

PABPC1 is usually considered the main source of cytoplasmic poly(A)-binding proteins in cells; however, the analysis indicates that some tissues have specifically produced PABP paralogs, which may act together with PABPC1 or partially substitute it. Not all of them were represented in proteomic datasets, but the available information, together with transcriptomic data, is sufficient to draw the following conclusions.

The level of the *PABPC1* transcript is relatively high in reproductive tissues (Supplementary Figure 3 A), while reduced in brain, heart, and skeletal muscles. The expression pattern is reflected in the INITIATION and, to some extent, TERMINATION+ complexes (Supplementary Figure 3B). Analysis of the *PABPC1* mRNA abundance among all PABPC paralogs (hereafter called “PABPC paralogs”) also revealed a significant decline of its expression in muscles (Figure 3A), suggesting that its role in this organ may be of lower importance.

**Figure 3.**
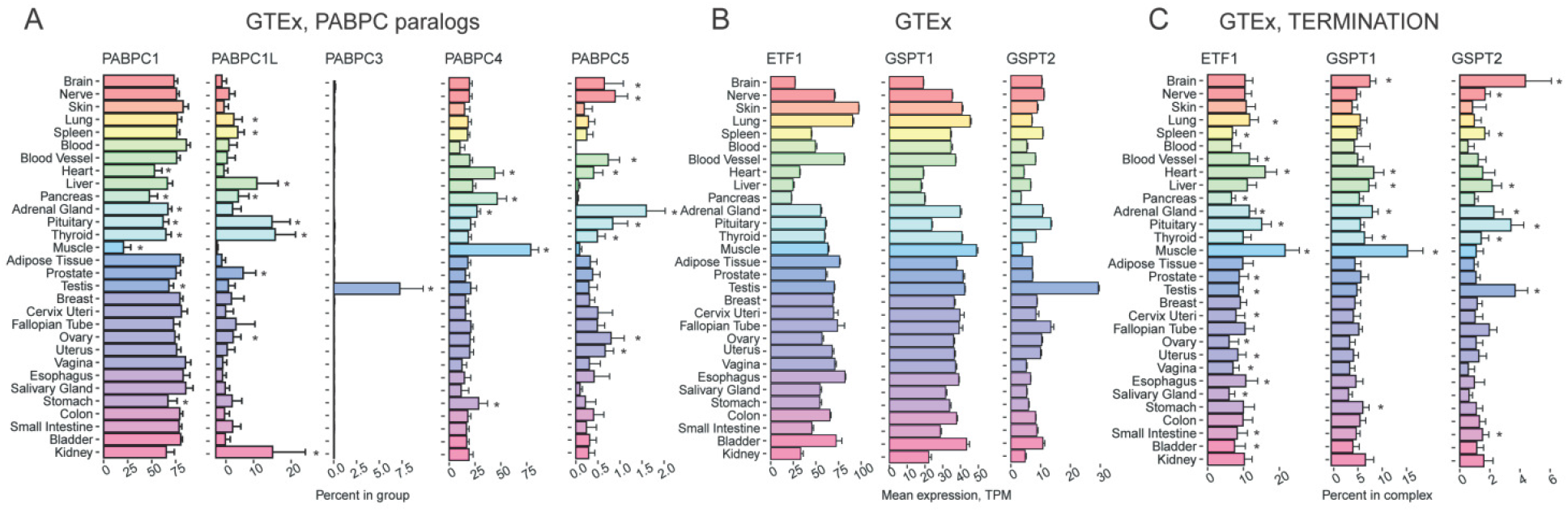
Tissue-specific expression pattern of genes encoding PABPC homologs and translation termination factors. **(A)** Percentage of *PABPC1, PABPC1L, PABPC3, PABPC4*, and *PABPC5* expression among the genes from the “PABPC” complex across various human tissues, according to GTEx. **(B)** Expression of the *ETF1, GSPT1*, and *GSPT1* genes encoding eRF1, eRF3a/GSPT1, and eRF3b/GSPT2 correspondingly in various human tissues according to GTEx. **(C)** Percentage of *ETF1, GSPT1*, and *GSPT1* expression among the genes from the “TERMINATION” complex across various human tissues, according to GTEx. TPM, Transcripts Per Kilobase Million; *, FDR corrected p-value < 0.01 in enrichment analysis (fgsea R package).

*PABPC1L*/*ePAB* was previously reported to have a gonad- and embryo-restricted expression [26,27]. This is consistent with its role in oocyte maturation in mice and its importance for female fertility in both mice and humans [65,66]. However, our analysis revealed also a high level of this transcript in pituitary and thyroid glands, as well as in liver and kidney, relative to that of other components of the “PABPCs complex” (Figure 3A). Its putative function in hormone-producing organs can contribute to its role in fertility.

PABPC3 was previously shown to be abundant only in testis [26]. Our analysis confirmed this highly specific expression pattern at the mRNA level (Figure 3A), but the protein level was unexpectedly high not only in testis, but also in adipose tissue, colon, and prostate (Supplementary Figure 3C). This case highlights the importance of addressing gene expression regulation at different levels. Importantly, *PABPC1*, *PABPC1L*, and *PABPC3* expression is perturbed in infertile men [67].

PABPC4 is able to compensate for the loss of PABPC1 [68]. In accordance with this, our approach revealed that *PABPC4* is upregulated at the mRNA and protein levels in muscle, heart, and pancreas in “PABPCs complex”, where *PABPC1* and *PABPC5* are downregulated (Figure 3A).

PABPC5 is an X-linked poly(A)-binding protein [69]. The analysis of its relative abundance within “PABPCs complex” revealed a relatively decreased level of the *PABPC5* transcript in pancreas, muscle, liver, salivary gland, and blood (Figure 3A, Supplementary Figure 3D). Interestingly, pancreas, muscle, and liver are among the most translationally active tissues containing the highest amount of ribosomes [44], so this pattern suggests its regulatory role in general protein synthesis.

eRF1 is an essential translation termination factor encoded by the *ETF1* gene. *ETF1* was claimed to be expressed at a high level in testis, brain, heart, and kidney and downregulated in the liver and colon [32]. The data obtained from GTEx (Figure 3B) only partially confirmed this, indicating the *ETF1* mRNA abundance is lower in liver, pancreas, kidney, and heart (likely the most metabolically active tissues [70]), as well as in brain, while there no significant difference between the colon and other tissues was observed. Remarkably, *ETF1*, while encoding a general translation factor, demonstrated an uneven transcription level within “TERMINATION+” group: its expression is higher in muscle, heart, and pituitary (Figure 3C).

*GSPT1* and *GSPT2* code for GTP-binding termination factors, eRF3a/GSPT1 and eRF3b/GSPT2, respectively [35]. The factors have at least partially redundant functions, as eRF3b can compensate for a silenced eRF3a [36]. We found that the level of *GSPT1* mRNA quite closely corresponds to that of mRNA encoding eRF1 (Figure 3B,3C). On the contrary, analysis of the *GSPT2* mRNA abundance suggests its specific relative increase in brain and testis, in comparison to other “TERMINATION” components (Figure 3C). The elevated *GSPT2* mRNA level in brain is consistent with Hoshino results [35] and may reflect a substituting role of eRF3b in nonproliferating cells, where eRF3a should be depleted.

GTPBP1 and GTPBP2 are GTPases involved in translational control: GTPBP1 has eEF1A-like elongation activity while GTPBP2 is likely involved in stalling ribosome rescue [71,72]. Their expression in mammalian tissues is poorly studied. Mouse brain was shown to have an elevated level of the *GTPBP1* transcripts [73], while testis and thymus are the organs with an increased abundance of the *GTPBP2* mRNA [74]. Our analysis of GTEx data revealed blood as a tissue with the highest level of the *GTPBP1* mRNA in the human body (Figure 4A), suggesting their specific role in blood cell differentiation and physiology. Unfortunately, these tissues were not represented in the proteomics data that we analyzed in this study.

**Figure 4.**
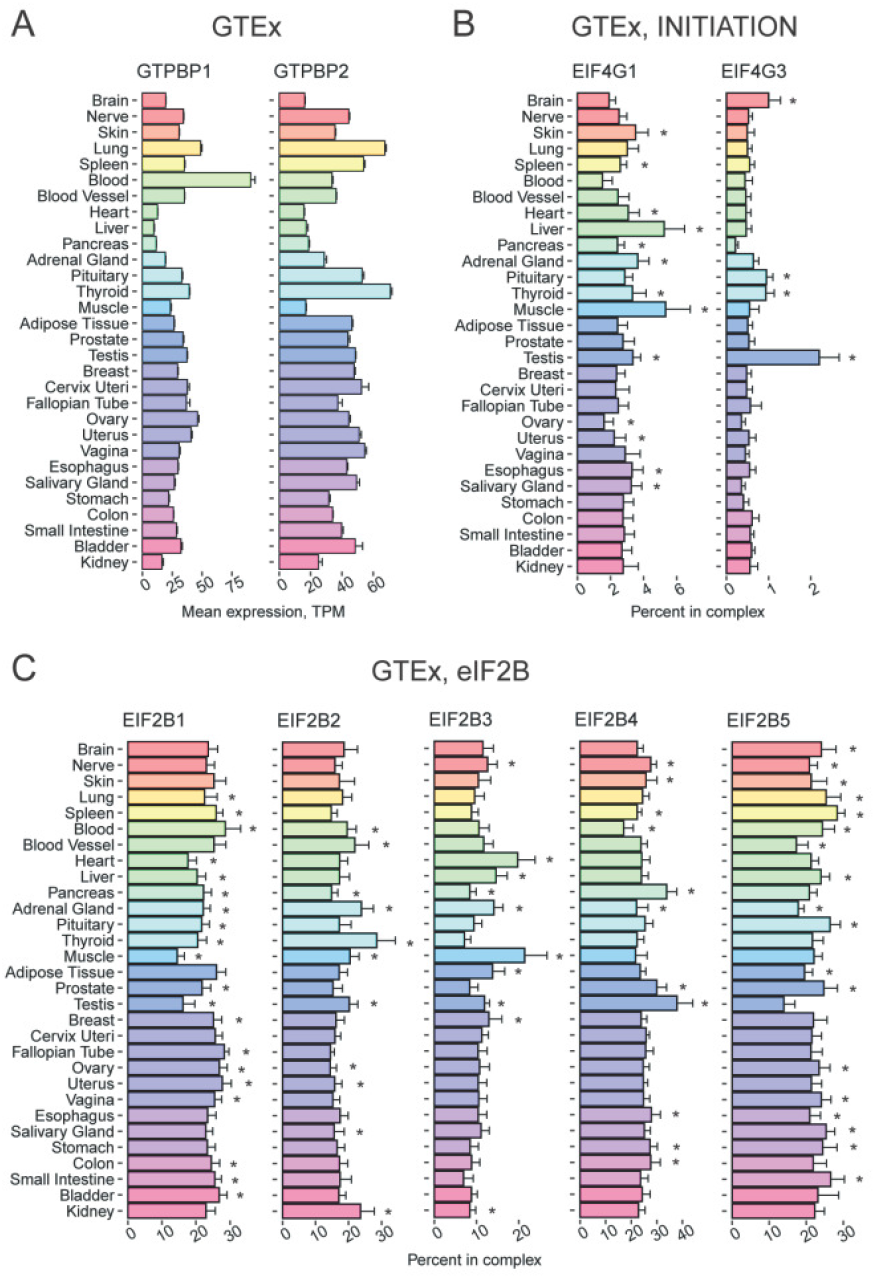
Tissue-specific expression pattern of genes encoding GTPBP1 and GTPBP2 proteins and eIF4G and eIF2B homologs. **(A)** Expression of the *GTPBP1* and *GTPBP2* genes in various human tissues according to GTEx. **(B)** Percentage of *EIF4G1* and *EIF4G3* among the genes from the “INITIATION” complex across various human tissues, according to GTEx. **(C)** Percentage of *EIF2B1*, *EIF2B2*, *EIF2B3*, *EIF2B4*, and *EIF2B5* among the genes from the “eIF2B” complex across various human tissues, according to GTEx. TPM, Transcripts Per Kilobase Million; *, FDR corrected p-value < 0.01 in enrichment analysis (fgsea R package).

*EIF4G3* codes for a paralog of eIF4G1. Both eIF4G1 and eIF4G3 are able to bind cap-binding protein eIF4E and RNA helicase eIF4A thus forming the trimeric translation initiation factor eIF4F. In humans, these paralogs share less than 50% identity and could have slightly different functions [75]. It was demonstrated previously that *EIF4G1* is highly expressed in liver and testis, while *EIF4G3* is upregulated in testis and fetal brain, and downregulated in lung, heart, liver, and placenta [76,77]. In agreement with these data, we found the level of *EIF4G3* transcript is elevated in testis (Supplementary Figure 4A), or in testis and brain if considered within the “INITIATION” group (Figure 4B). This is consistent with the fact that *EIF4G3* mutations cause male infertility [78]. Proteomic data confirmed the elevated level of eIF4G3 in testis and additionally revealed its elevated abundance in small intestine and smooth muscle (Supplementary Figure 4B). Thus, while eIF4G1 likely represents the major source of eIF4G activity in most tissues (Figure 4B, Supplementary Figure 4B), eIF4G3 might differentially contribute to translation in different organs, providing a fine-tuning of cap-dependent and alternative translation initiation pathways [79].

eIF2B is a guanine nucleotide exchange factor for eIF2. It consists of 5 subunits, α to ε, encoded by *EIF2B1*-*EIF2B5* genes, all of which are linked to a severe inherited human neurodegenerative disorder called Leukoencephalopathy with Vanishing White Matter, or VWM (for review, see [80]). Intriguingly, eIF2B subunits are known to form a number of differentially composed subcomplexes with non-identical activity [80]. Thus, we performed a comprehensive analysis of the relative abundance of individual eIF2B subunits and their mRNAs in comparison to those of all the subunits («EIF2B» group). At the mRNA level, we revealed the ubiquitous expression of five genes with no more than ~2-fold difference between tissues (Figure 4C), with probably some deviations in testis (with a lower *EIF2B1* and *EIF2B5* levels and a higher *EIF2B4* one), muscles and heart (lower *EIF2B1* and higher *EIF2B3* levels). Importantly, we observed no prominent specificity in subunit abundance either at mRNA or protein level in brain, suggesting that neural manifestation of VWM is not related to a distinct eIF2B composition in this organ. Mass-spectrometry data showed a more uneven subunit distribution (Supplementary Figure 4C), with putatively distinct stoichiometry in organs: for example, in heart, liver, thyroid, salivary gland, small intestine, and kidney EIF2B4 and EIF2B5 have an abundance above average levels, while EIF2B1 – below the average; in contrast, in gallbladder, pancreas, endometrium, and esophagus the proportions seems to be opposite. While the discrepancy at the mRNA and protein levels can be partially explained by a lower quality of the mass-spectrometry data, it should be noted that the abundance of eIF2B subunits is known to be regulated at the post-transcriptional level [81]. Overall, these findings suggest a diverse composition of the eIF2B complex across the human tissues.

### 3.4 New striking examples of tissue-specific expression of genes encoding translation factors

Translation initiation factor eIF4E1B belongs to the eIF4E cap-binding protein family, but it has a low affinity for the m^7^G-cap [82]. In amphibians, the eIF4E1B orthologue is part of a multisubunit complex that specifically inhibits the translation of some mRNAs in oocytes (Kubacka, 2015 #80;Minshall, 2007 #81}. Thus, it has been proposed that *EIF4E1B* expression is limited to ovaries and oocytes, where the factor plays a role in the translational repression of maternal mRNAs, while its canonical ortholog *EIF4E1* is transcribed ubiquitously to support the cap-dependent translation [82]. Unexpectedly, besides ovary, we found a high level of the *EIF4E1B* expression also in testis, retina, spinal cord, and brain, including pineal and pituitary glands (Figure 5A, Supplementary Figure 5A). This peculiar pattern of expression is probably related to that well documented for its putative partner CPEB [83,84].

**Figure 5.**
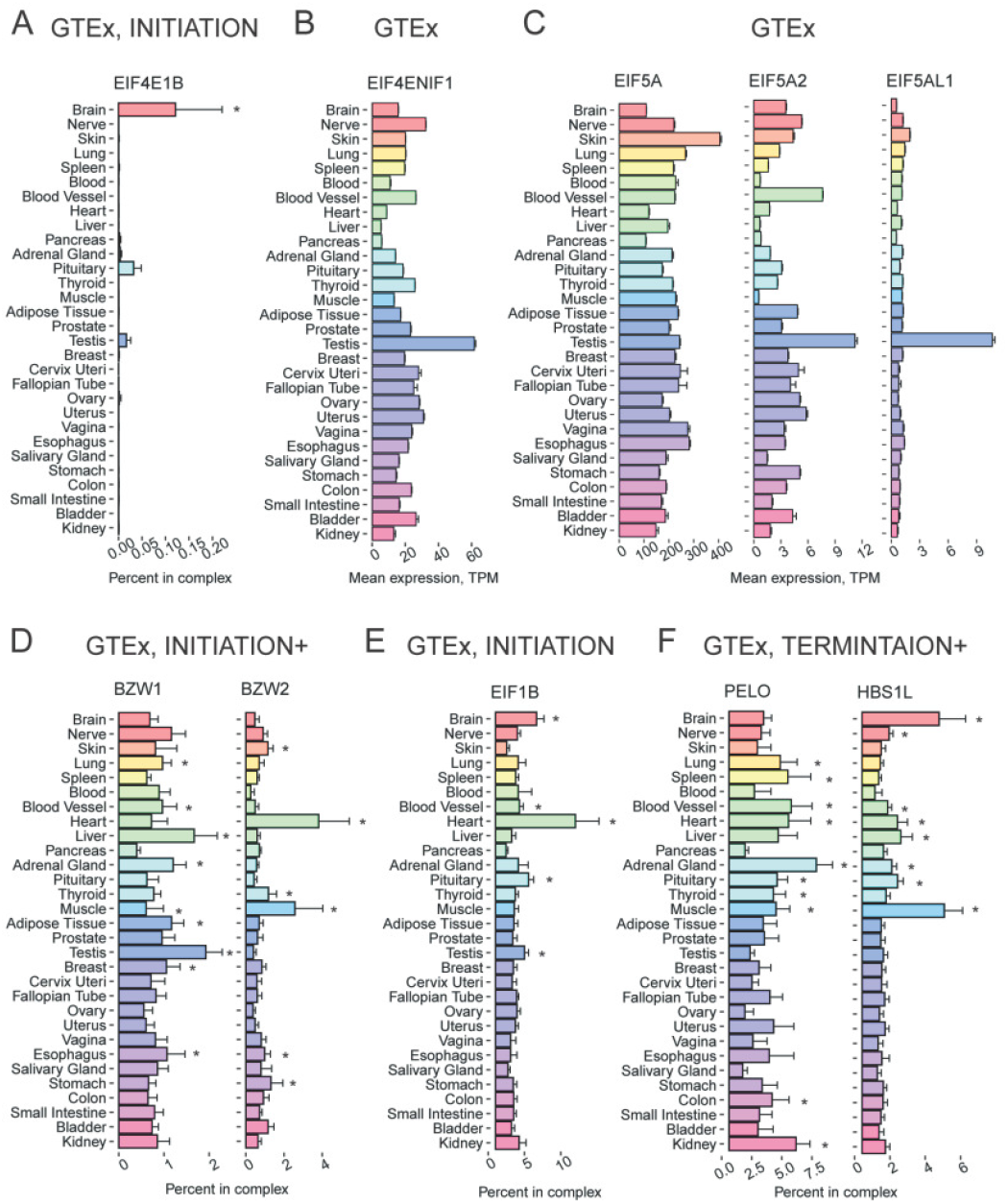
Tissue-specific expression pattern of translation-associated genes showing high tissue specificity. **(A)** Percentage of *EIF4E1B* expression among the genes from the “INITIATION” complex across various human tissues, according to GTEx. **(B)** Expression of the *EIF4ENIF1* in various human tissues according to GTEx. **(C)** Expression of the *EIF5A*, *EIF5A2*, and *EIF5AL1* in various human tissues according to GTEx. **(D)** Percentage of *BZW1* and *BZW2* expression among the genes from the “INITIATION+” complex across various human tissues, according to GTEx. **(E)** Percentage of *EIF1B* expression among the genes from the “INITIATION” complex across various human tissues, according to GTEx. **(F)** Percentage of *PELO* and *HBS1L* expression among the genes from the “TERMINATION+” complex across various human tissues, according to GTEx. TPM, Transcripts Per Kilobase Million; *, FDR corrected p-value < 0.01 in enrichment analysis (fgsea R package).

*EIF4ENIF1* encodes a multifunctional eIF4E-binding protein 4E-T that plays a role in the nucleocytoplasmic shuttling of eIF4E and in translational repression [85,86]. Interestingly, it binds not only the canonical eIF4E protein but also its orthologs (including eIF4E1B). Our analysis revealed that testis has the highest level of *EIF4ENIF1* transcript and protein (Figure 5B, Supplementary Figure 5B, C).

eIF5A, a translation elongation factor having the unique post-translation modification hypusine, is encoded in the human genome by three genes: *EIF5A, EIF5A2, EIF5AL1*. eIF5A1 (encoded by *EIF5A*) and EIF5AL1 are almost identical (98%), while eIF5A2 is slightly more diverged (84% identity) [29]. eIF5A1 is thought to be ubiquitous, while the *eIF5A2* expression has been reported to be restricted by brain and testis only, where it is represented by two or more isoforms with different 3’ UTR length [29,87].

Our analysis of GTEx and mass-spectrometry data confirmed the ubiquitous expression of *EIF5A*, but revealed a more complex pattern of the *EIF5A2* expression (Figure 5C, Supplementary figure 5D). In particular, eIF5A2 relative abundance is much lower at both mRNA and protein levels in liver and probably some other visceral organs but noticeably higher in testis. Additionally, we revealed a strict testis-specific expression of *EIF5AL1*: according to GTEx, this gene is almost exclusively expressed in this organ (Figure 5C). Yet, there was no information about the protein level of EIF5AL1. Taking into account the high similarity of eIF5A paralogs, it can be assumed that protein synthesis in male gonads specifically requires a higher eIF5A concentration, although a distinct testis-specific function of eIF5AL1 cannot be ruled out either.

BZW1/5MP2 and BZW2/5MP1 are eIF5-mimic proteins that regulate the stringency of start site selection [88]. The relative level of *BZW1* mRNA varies moderately between tissues, being highest in liver and testis, and lowest in pancreas (Figure 5D). In contrast, the relative abundance of *BZW2* transcript (which is overall lower if assessed in the context of “INITIATION+” group) is noticeably high in only two tissues, heart and muscle (Figure 5D). The elevated expression of *BZW2* in heart is also evident at the protein level, and it is also high in placenta (Supplementary Figure 5E). This suggests that BZW1/5MP2 is likely a general regulator of protein synthesis, while BZW2/5MP1 could be a specialized factor for tissue- and mRNA-specific translational control.

A poorly studied protein eIF1B is 92% identical to its paralog eIF1, which plays a major role in the start codon selection along with eIF5[89]. While *EIF1* relative expression is ubiquitous with little variations across organs (Figure 5E), *EIF1B* mRNA relative abundance is clearly higher in heart and brain, while lower in skin and pancreas (Figure 5E). An even more complex distribution of eIF1B across tissues is observed at the protein level (Supplementary Figure 5F), in agreement with a proposed regulation at the level of translation [90]. The high similarity between eIF1B and eIF1, as well as some indirect data (see [89] and references therein), suggests the functional redundancy of these proteins. Thus, an increased total amount of the eIF1/eIF1B activity could contribute to tissuespecific regulation of initiator codon selection similar to that observed in cells artificially overexpressing eIF1 [91].

We also detected tissue-specific expression of some translation quality control factors that had not been reported before. *PELO* and *HBS1L* encode non-canonical termination factors mediating the dissociation of inactive, vacant, or stalled ribosomes (for review, see [92]). At least one case of a cell-type specific translational control by PELO and HBS1L abundance has been reported [93]. It is also important to note that HBS1L deficiency causes organ-specific defects during development [94]. These facts suggest the involvement of the factors in tissue-specific regulation of gene expression. Although PELO and HBS1L are thought to work in tandem, analyses of transcriptomic and massspectrometry data suggest that their abundances do not correlate well across tissues. In particular, we would like to report the inverse ratio of the encoding mRNAs in brain and muscle (where the relative level of *HBS1L* mRNA is higher than that of *PELO*), vs. adrenal gland, kidney, and spleen (where the proportion is inverse), see Figure 5F. At the protein level, liver, gallbladder, duodenum, and testis have relatively higher levels of HBS1L than PELO, while thyroid, prostate, and bladder demonstrate the opposite tendency (Supplementary Figure 5G).

### 3.5 Genes encoding some ARSases also have a pronounced tissue-specific expression

Aminoacyl-tRNA synthetases (ARSases) are key components of protein synthesis machinery. Surprisingly, we found that some of them also have diverse expression patterns across organs and tissues. As before, we analyzed absolute and relative mRNA and protein abundance of ARSases within functional groups. In Metazoa, some ARSases form a multiprotein complex with auxiliary factors [95]. Thus, we used two functional groups: “ARSases” and “ARSase COMPLEX” (Table 1).

*HARS1* gene encoding histidyl-tRNA synthetase (*HARS1*) has been previously reported as highly expressed in heart, brain, liver, and kidney [96]. According to our analysis, the corresponding mRNA is relatively more abundant in brain, especially in pituitary, as compared to other ARSases, although more or less homogenously distributed in other organs (Figure 6A, Supplementary Figure 6A). The *TARSL2/TARS3* gene encoding threonyl-tRNA synthetase is relatively upregulated in brain, spinal cord, muscle, and heart. The data from GTEx and FANTOM revealed that *TARSL2* was efficiently transcribed in brain and heart (Figure 6A, Supplementary Figure 6A). It is consistent with its high abundance in muscle and heart in mice, as reported earlier [97]. Lung, blood, and placenta are the tissues with increased relative levels of tryptophanyl-tRNA synthetase (*WARS*) transcript and protein (Figure 6B, Supplementary Figure 6B).*AIMP2* codes for a scaffold protein of the Multiple ARSase Complex (MARS), AIMP2, which has 2 partners: AIMP1 and EEFE1/AIMP3 [95]. The data obtained from all three databases indicated that the relative abundance of *AIMP2* transcript (as compared to that of all “ARSase COMPLEX” components, see Table 1) is elevated in muscle tissues and testis (Figure 6C). In contrast, *AIMP1* mRNA and protein did not show any prominent tissuespecific pattern (Figure 6C and Supplementary Figure 6C), The third key MARS component, EEF1E1, shows no unambiguous correlation between mRNA and protein levels, suggesting regulation of its abundance at the translational level.

**Figure 6.**
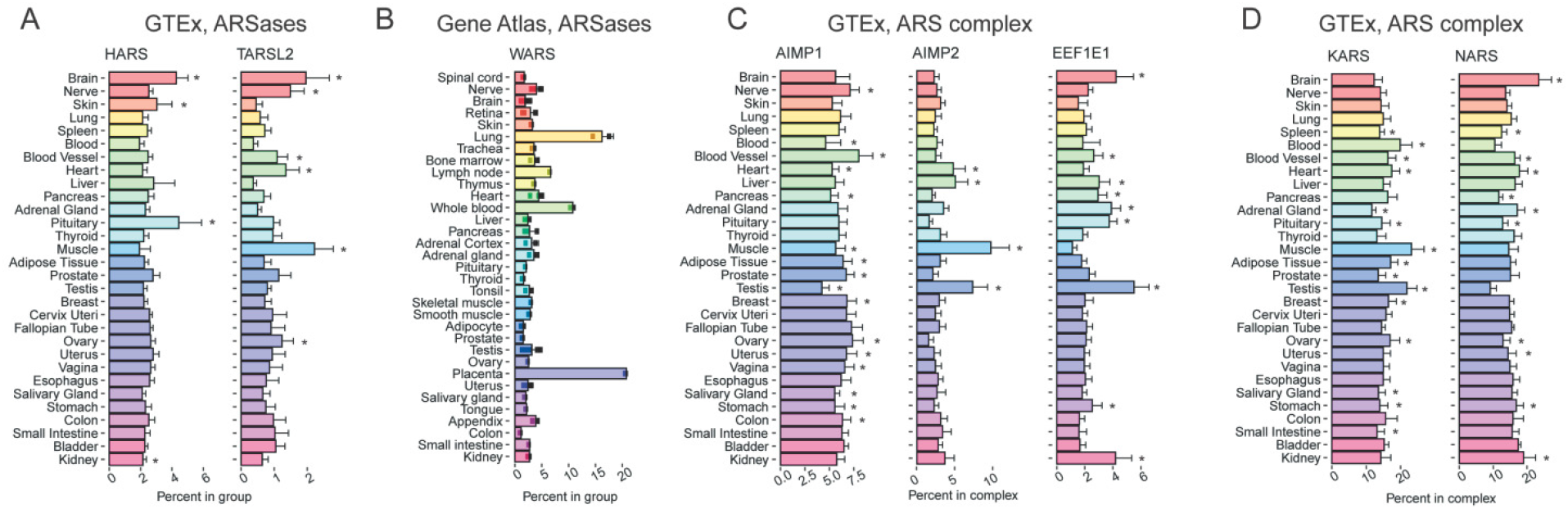
Tissue-specific expression pattern of genes encoding several aminoacyl-tRNA-synthetases (ARSases) showing tissue specificity. **(A)** Percentage of *HARS* and *TARSL2* expression among the genes from the “ARSs” complex across various human tissues, according to GTEx. **(B)** Percentage of *WARS* expression among the genes from the “ARSs” complex across various human tissues, according to Gene Atlas [46]. **(C)** Percentage of *AIMP1, AIMP2*, and *EEF1E1* expression among the genes from the “ARS complex” complex across various human tissues, according to GTEx. **(D)** Percentage of *KARS* and *NARS* expression among the genes from the “ARS complex” complex across various human tissues, according to GTEx. *, FDR corrected p-value < 0.01 in enrichment analysis (fgsea R package).

ARSase components of MARS can also be differentially distributed across tissues. A more or less homogenous expression is exemplified by *KARS* gene coding for lysyl-tRNA synthetase (Figure 6D and Supplementary Figure 6D). In contrast, the relative expression of *NARS* (encoding asparaginyl-tRNA synthetase), although similarly high in most tissues, is further elevated in brain but repressed in testis, as evident at both mRNA and protein levels (Figure 6D and Supplementary Figure 6D). Thus, MARS composition may vary in different human organs and tissues.

## 4 Discussion

mRNA translation is one of the basic processes in a living cell. However, it is clear that in a multicellular organism, various cell types differentially rely on the synthesis of new proteins and thus require different amounts of translation machinery components. In addition, the composition of this apparatus can contribute to the translational control of gene expression. Thus, human tissues and organs may have diverse concentrations, ratios, and compositions of ribosomes, tRNAs, translation factors, and auxiliary proteins. Differential abundance of tRNA species in human tissues was reported previously [22,23]. Expression levels of genes coding for ribosomal proteins were reported to vary significantly across human tissues as well [20,98]. In 2012, Maria Barna and colleagues proposed a concept of the specialized ribosome [99], stating that different mRNA species can be translated by heterogeneously composed ribosomes, depending on cell types and conditions. The differential composition of ribosomes was indeed confirmed in a number of subsequent studies [19–21].

Here we present evidence that many translation factors, both general and non-canonical ones, as well as auxiliary proteins like ARSases, also show tissue-specific expression patterns, dictating diverse composition of translation apparatus in different cell types. In particular, we systematically explored three transcriptomic databases (GTEx, FANTOM, and Gene Atlas) and one source of proteomic data [1], and found a number of translation machinery components with strict tissuespecific appearance: eEF1A1, eEF1A2, PABPC1L, PABPC3, eIF1B, eIF4E1B, eIF4ENIF1, and eIF5AL1; we then identified a differential relative abundance of PABP and eRF3 paralogs, eIF2B subunits, eIF5-mimic proteins, and some ARSases, suggesting that even some general factors may have tissue-specific functions; we also noted sexual dimorphism in the repertoire of translation-associated proteins encoded in sex chromosomes: eIF1A, eIF2γ, and DDX3.

Recently, a global analysis of tRNA and translation factor expression in human cancers was performed [100]. The authors revealed overall overexpression of tRNA modification enzymes, ARSases, and translation factors, which may play a role in the activation of protein synthesis across multiple cancer types. However, cancer cells are well known to have deregulated gene expression, so it was very important to show that normal human tissues can have a diverse composition of translation factors as well.

Obviously, normal mammalian tissues have highly variable levels of protein synthesis, depending on metabolic rate, secretory activity, and cell proliferation status. A number of studies assessed protein synthesis rates across tissues by metabolic labeling [41–43]. These analyses revealed the highest amino acid incorporation and/or protein turnover in small intestine and pancreas, intermediate in kidney, spleen, and liver; and the lowest in lung, heart, brain, muscle, and adipose tissue. It was also shown that the protein synthesis rate is tightly coupled to cell metabolic fluxes [101]. Metabolic activities of mammalian tissues vary significantly in a row from heart and kidney (the most active ones) to skeletal muscle and adipose tissue [70]. Ribosome amounts also vary dramatically (~50-fold) between the tissues, with more ribosomes present in pancreas, salivary gland, liver, and intestine; medium in muscle, ovary, kidney, and brain cortex; and lower in thymus, spleen, heart, lung, and cerebellum [44]. All this makes it clear that for the assessment of tissue-specific features of translation machinery, one should not use absolute values of mRNA or protein abundances of its components. Thus, in our analyses, we use a novel methodology when we combine translation factors into functional groups and calculate the relative mRNA or protein abundance of the particular component within the groups.

This approach allowed comparing the relative concentration of translation factors and associated proteins and revealed some tissue-specific peculiarities in the composition of protein synthesis machinery contributing to translational landscapes of human tissues. For example, we would like to note a potentially higher total activity of eIF1, eIF1A, and BZW in heart due to an “additional source” of these factors, i.e. the elevated expression of *EIF1B*, *EIF1AX*, and *BZW2* genes specifically in this organ (Figures 2A,5D-E). These factors are known to enhance the stringency of start site selection [45,88] and thus should remodel a pattern of efficiently translated mRNAs. Future studies using ribosome profiling of animal organs [2,12–15] could shed some light on this issue.

Our study also draws attention to sexual dimorphism in the abundance of some translation machinery components. At least three important factors, eIF1A, eIF2γ, and DDX3, are encoded in sex chromosomes (by the X-linked *EIF1AX, DDX3X*, and *EIF2S3* genes, and Y-linked *EIF1AY* and *DDX3Y* genes). In addition to the predictable lack of the Y-linked gene expression in female organs, this issue clearly contributes to total amounts of translation factors in some organs (e.g. the higher eIF1A level in the heart due to a “double portion” of EIF1AX/Y expression, or the lower eIF2γ level in testis than in female organs due to *EIF2S3* X-inactivation escape, Figure 2).

Our analysis identified testis and brain as organs with the most diverged expression of translation-associated genes. Although it is well known that neurons and nervous tissues have peculiarities in protein synthesis [102,103], translation regulation in testis has not been extensively studied yet. A recent comparison of the mammalian transcriptome and proteome revealed that the correlation between the transcriptome and the translatome in testis is much lower than in the brain and liver [2]. It should also be noted that the brain and testis are immunologically privileged organs and this may bring some bias in transcriptomic and proteomic analysis of their content. It was also shown recently that testis and brain express genes that are enriched in rare codons both in humans and flies [104].

Recent studies show that different tissues can have different tRNA repertoires and codon usage [16,17]. The adaptation of the tRNA pool was found to be largely related to a tissue proliferative state [16]. Then, two clusters of tissues with an opposite pattern of codon preferences were identified [105]: some tissues (including kidney, muscle, heart, liver, colon, fat, and ovary) generally favor C/G-ending codons, while others (including lung, brain, pancreas, spleen, small intestine, adrenal and salivary glands, placenta, and testis) better tolerate translation of rare A/T-ending codons. Although we were unable to find any obvious correlation between these clusters and the distribution of elongation factors or ARSases, more in-depth analysis may reveal such patterns in future.

Finally, we would like to emphasize the importance of using both transcriptomic and proteomic analyses. Although mRNA levels are primary determinants for protein abundance (for review, see [106]), translation regulation can significantly contribute to the levels of translation machinery components [90]. Unfortunately, for many poorly expressed genes, proteomic data are either lacking or statistically unreliable, while transcriptomic datasets from GTEx, FANTOM, and Gene Atlas are usually enriched in reliable information about most human genes. The discrepancy between the transcript and protein levels that we reported for some translation-associated genes can be a starting point for the investigation of their post-transcription regulation.

## 5 Conclusions

In summary, in this study we systematically explored the tissue-specific expression of translation-associated genes and found new cases of differentially represented translation factors and auxiliary proteins in human organs and tissues. These findings contribute to our understanding of translational control in health and disease.

## 6 Author Contributions

Conceptualization, S.E.D., A.S.A., and I.V.K.; experimental design, A.S.A., I.V.K., A.A.E., and S.E.D.; methodology, A.S.A., I.V.K., A.A.E., and S.E.D.; data analysis, A.S.A.; discussion of results, A.S.A., N.M.K., N.E.M., A.A.E., I.V.K., and S.E.D.; writing of manuscript and editing, A.S.A., N.M.K., N.E.M., A.A.E., I.V.K., and S.E.D.; project administration, S.E.D.; funding acquisition, S.E.D. All authors have read and agreed to the published version of the manuscript.

## 7 Funding

This study was supported by the Russian Science Foundation: RSF grants no. 18-14-00291 to S.E.D.(conceptualization, design, and data analysis) and no. 20-74-10075 to I.V.K. (data processing pipeline).

## 8 Acknowledgments

N.M.K., N.E.M., A.A.E., and S.E.D. are members of the Interdisciplinary Scientific and Educational School of Moscow University «Molecular Technologies of the Living Systems and Synthetic Biology».

## 9 Conflicts of Interest

The authors declare that the research was conducted in the absence of any commercial or financial relationships that could be construed as a potential conflict of interest.

## Notes

### Competing Interest Statement

The authors have declared no competing interest.

