## Supplementary figures for "Human tissues exhibit diverse composition of translation machinery"

---

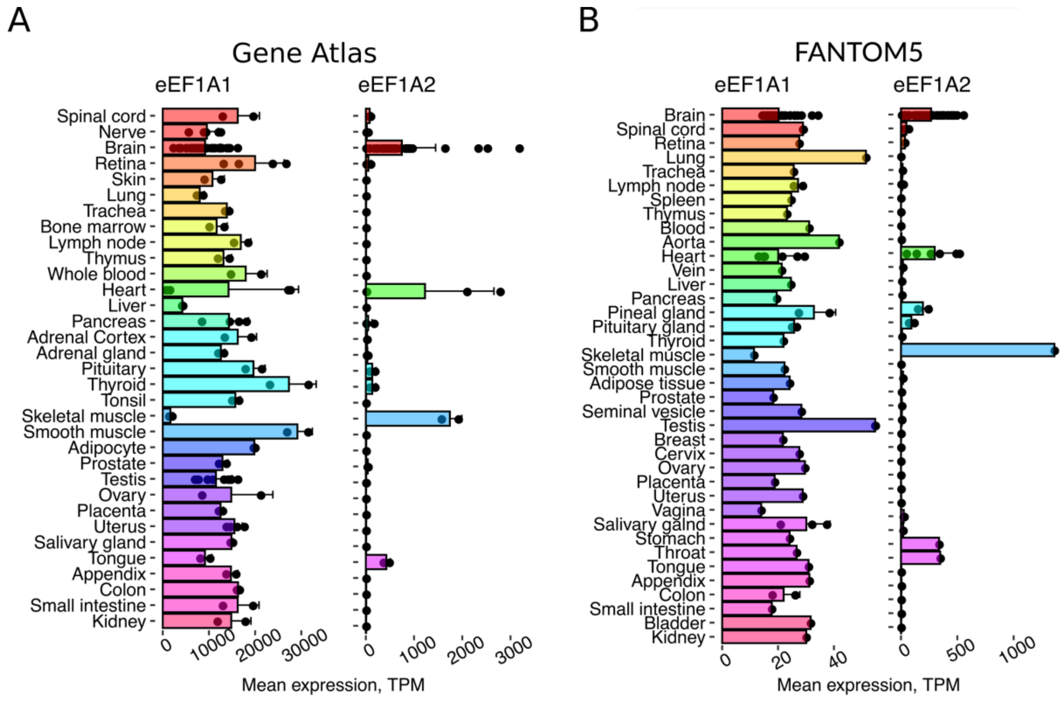


**Supplementary Figure 1.** Tissue-specific expression pattern of two human genes encoding eEF1A paralogs, *EEF1A1* and *EEF1A2*. **(A)** Expression of the *EEF1A1* and *EEF1A2* genes in various human tissues according to Gene Atlas [1]. **(B)** Expression of the *EEF1A1* and *EEF1A2* genes in various human tissues according to FANTOM5. TPM, Transcripts Per Kilobase Million.

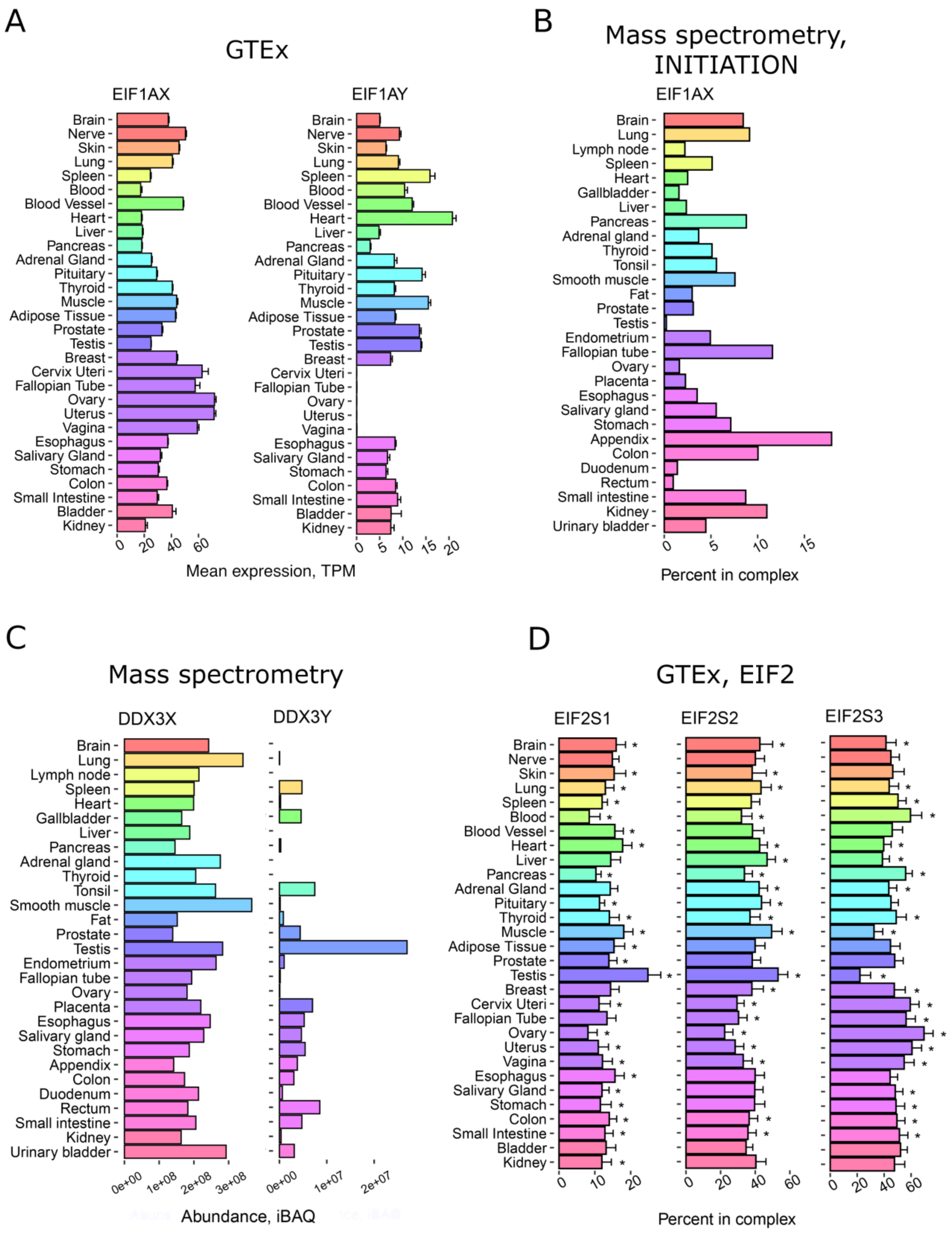


**Supplementary Figure 2.** Tissue-specific expression pattern of translation-associated genes localized in sex chromosomes. **(A)** Expression of the *EIF1AX* and *EIF1AY* genes in various human tissues according to GTEx. **(B)** Percentage of *EIF1AX* expression among the genes from the "INITIATION" complex across various human tissues, according to proteomic analysis [2]. **(C)** Levels of DDX3X and DDX3Y proteins in various human tissues according to proteomic analysis. **(D)** Percentage of *EIF2S1*, *EIF2S2*, and *EIF2S3* expression encoding α, β, and γ-subunits of eIF2 complex within the "EIF2" complex across various human tissues, according to GTEx. *, FDR corrected p-value < 0.01 in enrichment analysis (fgsea R package).

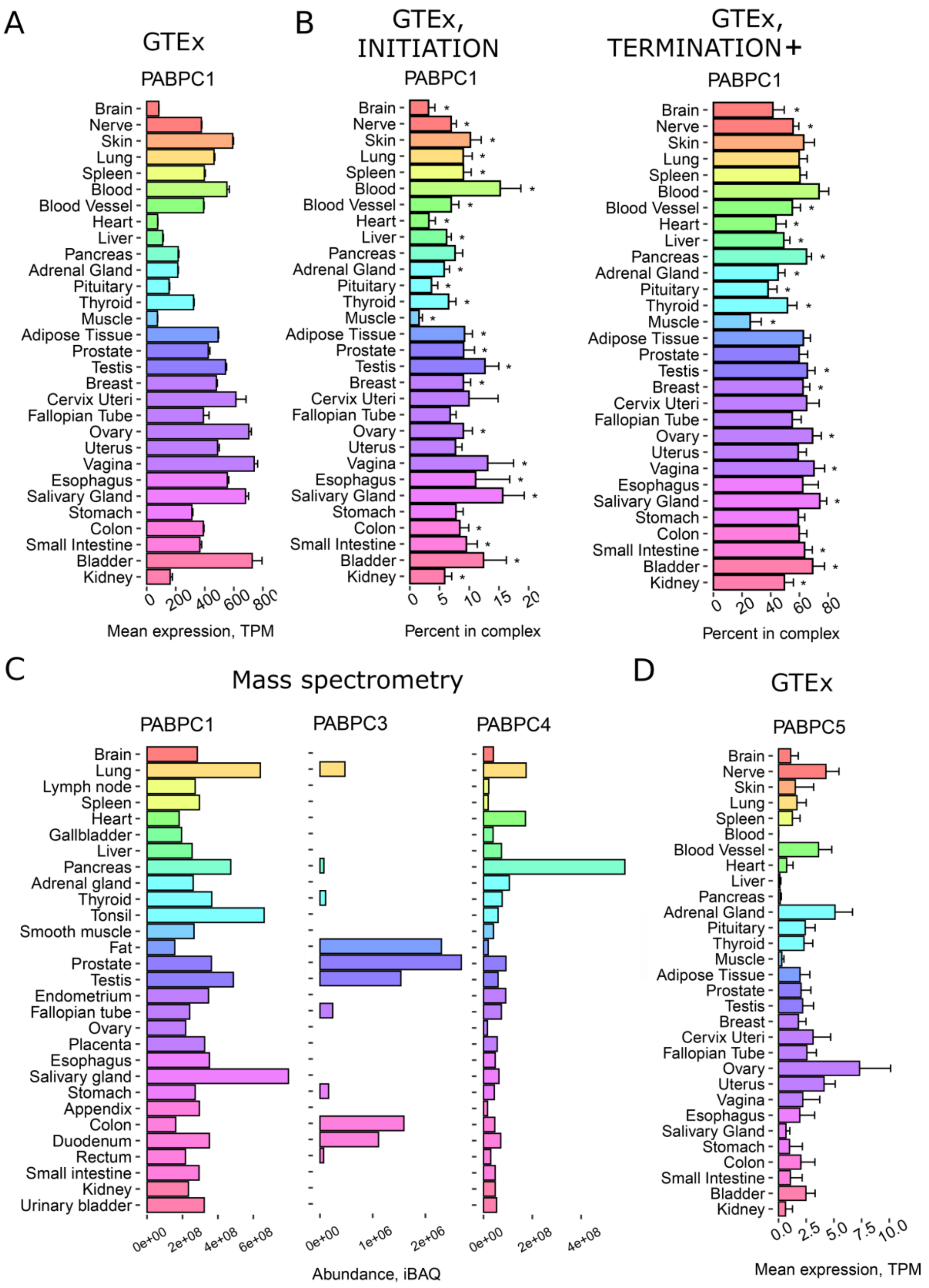


**Supplementary Figure 3.** Tissue-specific expression pattern of genes encoding PABPC homologs. **(A)** Expression of the *PABPC1* in various human tissues according to GTEx. **(B)** Percentage of *PABPC1* expression among the genes from the "INITIATION" and "TERMINATION+" complexes across various human tissues, according to GTEx. **(C)** Levels of PABPC1, PABPC3, and PABPC4 proteins in various human tissues according to proteomic analysis [2]. **(D)** Expression of the *PABPC5* in various human tissues according to GTEx. TPM, Transcripts Per Kilobase Million; *, FDR corrected p-value < 0.01 in enrichment analysis (fgsea R package).

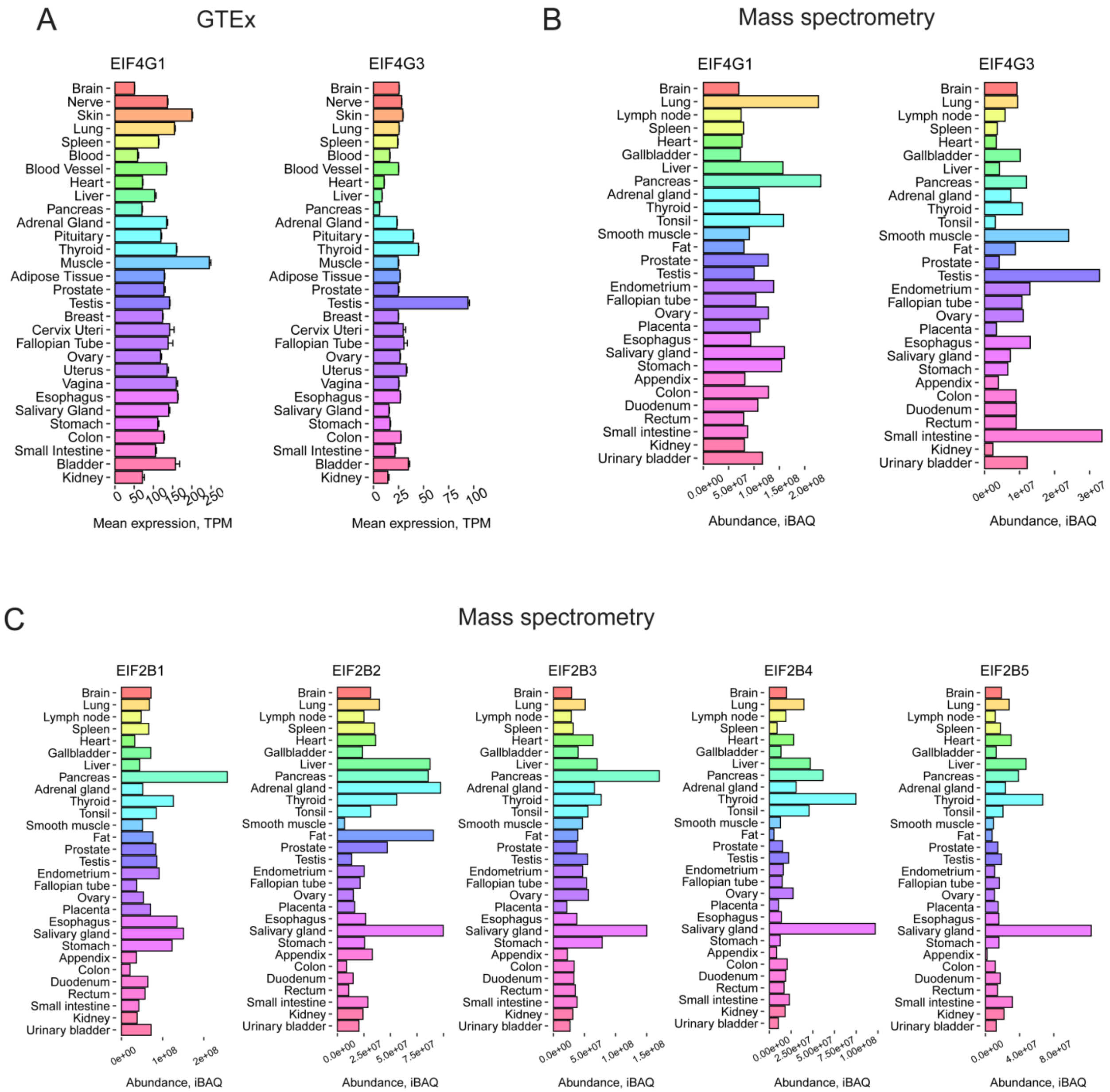


**Supplementary Figure 4.** Tissue-specific expression pattern of genes encoding GTPBP1 and GTPBP2 proteins and eIF4G and eIF2B homologs. **(A)** Expression of the *EIF4G1* and *EIF4G3* genes in various human tissues according to GTEx. **(B)** Levels of EIF4G1 and EIF4G3 proteins in various human tissues according to proteomic analysis [2]. **(C)** Levels of eIF2B homologs in various human tissues according to proteomic analysis. TPM, Transcripts Per Kilobase Million.

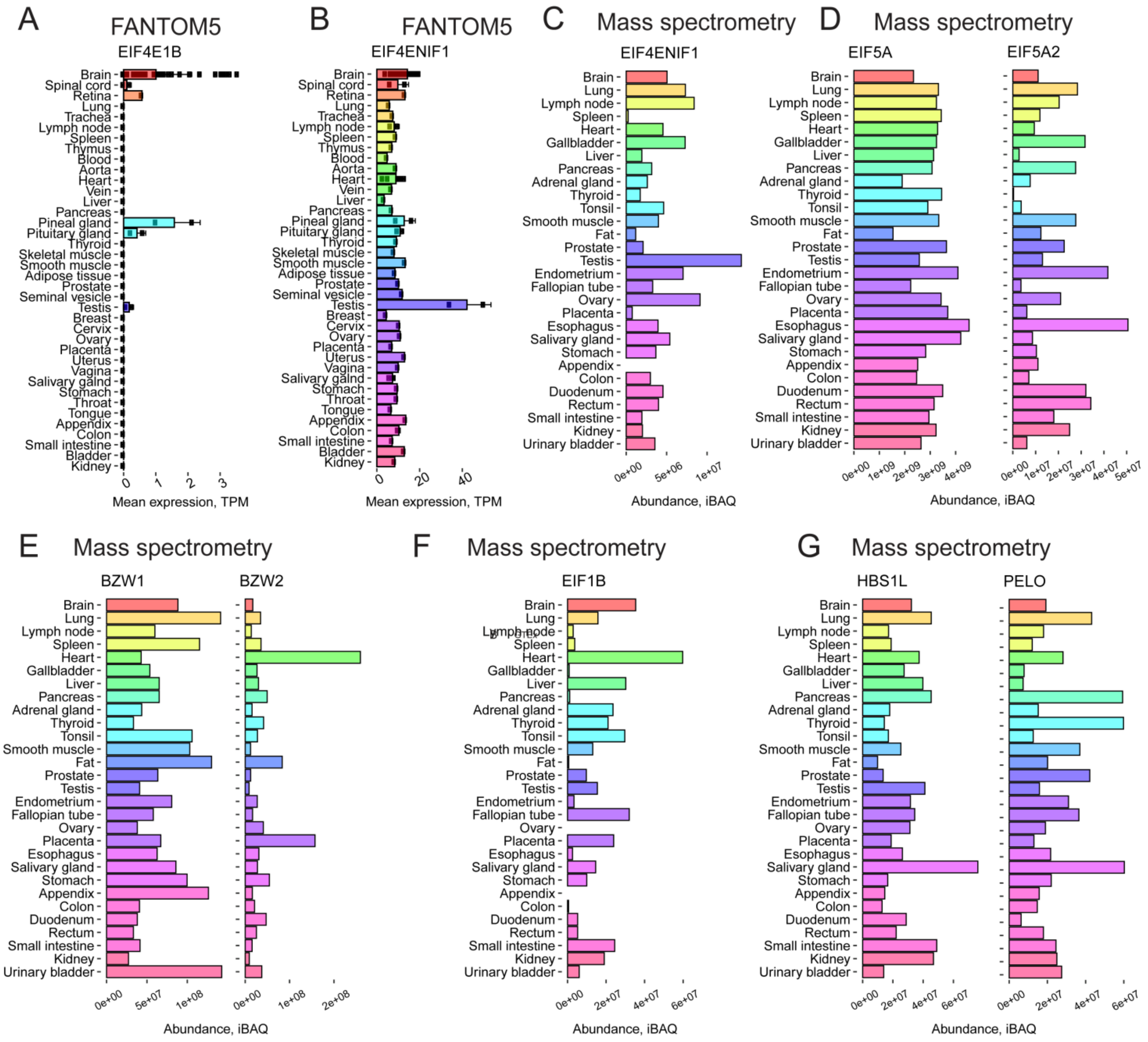


**Supplementary Figure 5.** Tissue-specific expression pattern of translation-associated genes showing high tissue specificity. **(A)** Expression of the *EIF4E1B* in various human tissues according to FANTOM5. **(B)** Expression of the *EIF4ENIF1* in various human tissues according to FANTOM5. **(C)** Levels of EIF4ENIF1 protein in various human tissues according to proteomic analysis [2]. **(D)** Levels of eIF5A homologs in various human tissues according to proteomic analysis. **(E)** Levels of BZW1/5MP2 and BZW2/5MP1 proteins in various human tissues according to proteomic analysis. **(F)** Levels of eIF1B protein in various human tissues according to proteomic analysis. **(G)** Levels of HBS1L and PELO proteins in various human tissues according to proteomic analysis. TPM, Transcripts Per Kilobase Million.

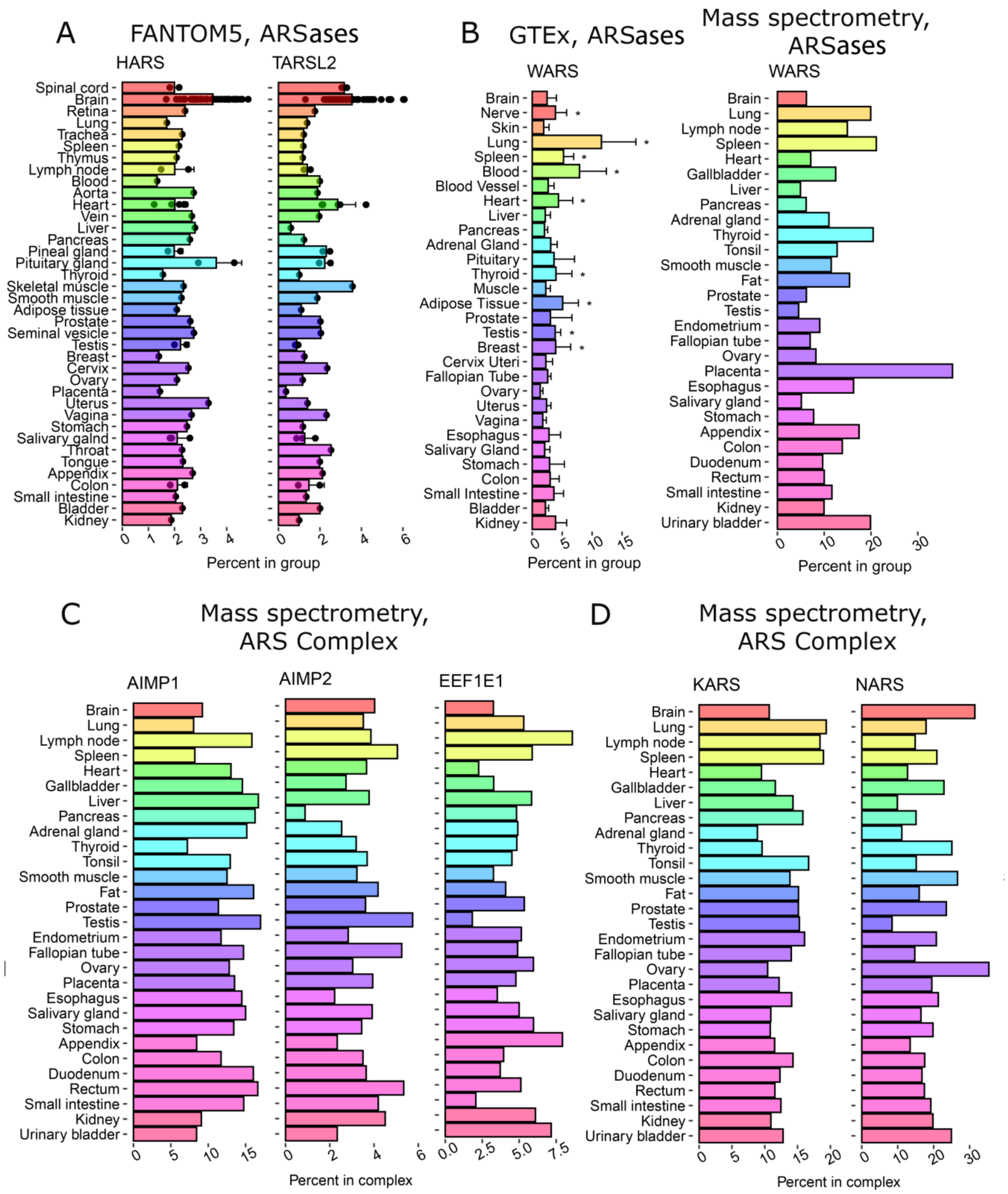


**Supplementary Figure 6.** Tissue-specific expression pattern of genes encoding several aminoacyl-tRNA-synthetases (ARSases) showing tissue specificity. **(A)** Percentage of *HARS* and *TARSL2* expression among the genes from the "ARSs" complex across various human tissues, according to FANTOM5. **(B)** Percentage of *WARS* expression among the genes from the "ARSs" complex across various human tissues, according to GTEx and proteomic analysis [2]. **(C)** Percentage of *AIMP1*, *AIMP2*, and *EEF1E1* expression among the genes from the "ARS complex" complex across various human tissues, according to proteomic analysis. **(D)** Percentage of *KARS* and *NARS* expression among the genes from the "ARS complex" complex across various human tissues, according to proteomic analysis. TPM, Transcripts Per Kilobase Million; *, FDR corrected p-value < 0.01 in enrichment analysis (fgsea R package).
